# Simulating human foot mechanics during walking based on an anatomically detailed forward dynamic finite element model

**DOI:** 10.1101/2025.10.12.681316

**Authors:** Kohta Ito, Yuka Matsumoto, Hiroyuki Seki, Takeo Nagura, Naomichi Ogihara

**Author notes:** These authors have contributed equally to this work and share first authorship (Kohta Ito, Yuka Matsumoto). Corresponding Author Naomichi Ogihara, Department of Biological Science, Graduate School of Science, The University of Tokyo, 7-3-1 Hongo, Bukyo-ku, Tokyo 113-0033, Japan.

## Abstract

**Purpose:** Forward dynamic musculoskeletal simulation is a powerful computational approach for investigating the biomechanics of human locomotion. However, existing models often oversimplify foot anatomy, thereby limiting our understanding of the role of detailed foot morphology in gait mechanics. In this study, we developed an anatomically accurate three-dimensional finite element (FE) model of the human foot to simulate its dynamic behavior during the stance phase of walking using an explicit forward dynamics approach.

**Methods:** The model incorporated detailed representations of bones, soft tissues, ligaments, and the plantar aponeurosis, and was driven by experimentally measured tibial kinematics and estimated muscle forces.

**Results:** Simulation results reasonably matched experimental data on ground reaction forces, plantar pressure distributions, and bone movements, confirming the model’s ability to replicate key aspects of foot-ground interactions during walking. Moreover, the model enabled the estimation of internal forces, stresses, and strains in foot structures that are not directly measurable in vivo, offering new insights into the biomechanics underlying foot pathologies.

**Conclusions:** This study potentially provides a robust framework for exploring the form–function relationship of the human foot, with applications in evolutionary biology, clinical interventions, and the study of locomotor disorders.

## Introduction

Forward dynamic musculoskeletal simulation is a computational technique to calculate muscle-driven skeletal movements through the integration of differential equations governing the dynamics of a musculoskeletal system [55, 37]. Extensive research in forward dynamic simulations has focused on bipedal walking, aiming to understand the complex biomechanics and neurocontrol mechanisms governing human locomotion [e.g., 50, 54, 35, 44, 21, 32, 2, 46, 10, 36]. These simulations also hold significance in predicting alterations in the kinematics, kinetics, and energetics of walking resulting from virtual alterations to the musculoskeletal system. For example, Oku et al. [36] demonstrated using a forward dynamic musculoskeletal simulation that the manipulation of the foot structure from digitigrade to humanlike plantigrade to allow heel contact results in improved cost of transport, suggesting that evolutionary changes in the foot structure were crucial for the acquisition of humanlike efficient bipedal locomotion. Such virtual alterations offer a unique opportunity for exploring the potential impact of anatomical changes or interventions on human gait, thus providing valuable insights into understanding form-function relationship of the musculoskeletal system and human movement disorders.

However, in these models, it is not feasible to assess how variations in the morphology of small foot bones impact the forces acting on the foot, and thus, the overall dynamics of the body. This limitation arises because the foot is typically represented as either a single rigid body or a two rigid-link model comprising the foot and toe. Such simplifications prevent accurate representation of the complex movements and forces within the foot. On the other hand, computer simulations of the complete human foot during walking, based on anatomically and physiologically accurate finite element (FE) models and loading conditions, offer a pathway to elucidate the form–function relationship of the human foot. These simulations have the potential to analyze and predict alterations in foot function resulting from virtual modifications to foot morphology. Consequently, they facilitate investigations into how differences in foot morphology affect the mechanical performance of bipedal locomotion.

The objective of this study is to develop a three-dimensional (3D) FE model of the human foot, grounded in anatomical accuracy, to simulate the dynamic behavior observed during the stance phase of bipedal walking using forward dynamics. While a previous study conducted a two-dimensional dynamic FE analysis of the foot during walking to elucidate its dynamic and functional interaction with the ground [39], more recent efforts have seen the emergence of 3D explicit dynamic FE analyses of the human foot during both walking [1, 52, 30] and running [7]. However, these 3D FE foot models were typically validated primarily against experimentally obtained plantar pressure distributions applied to the foot’s plantar surface, neglecting validation against horizontal (anteroposterior and mediolateral) ground reaction forces and bone kinematics. Consequently, while these studies successfully replicated foot mechanics during walking and running using 3D human foot models, there may exist potential limitations in accurately reproducing foot dynamics. The realization of a forward dynamic simulation of human walking, incorporating an anatomically realistic foot model that accurately replicates the true kinematics and kinetics of bipedal locomotion, holds promise as a powerful tool for elucidating the relationship between form and function in the musculoskeletal system and understanding human movement disorders.

## Method

### Human foot model

In the present study, the 3D FE model of the human foot developed in Ito et al. [20] was updated to be anatomically more realistic and accurate (Figure 1a). The foot model was constructed based on CT scan data of a healthy male adult (age, 42 years; height, 172 cm; weight, 72 kg; foot length, 25.5 cm) with no history of orthopaedic and neuromuscular impairments. The foot model consisted of a total of 23 bones: tibia, fibula, talus, calcaneus, navicular, cuboid, three cuneiforms, five metatarsals, proximal and distal hallucal phalanges, and four fused phalanges of the second to fifth rays, and three sesamoids, all meshed with tetrahedral elements. The bones were represented as an isotropic linear elastic material, and the Young’s modulus, Poisson’s ratio, and density were set to 7300 MPa, 0.3, and 0.0015 g/mm^3^, respectively [8, 39]. The encapsulated soft tissues between the outer foot surface and bone surface were also meshed with tetrahedral elements. The encapsulated soft tissue was defined as a hyperelastic Ogden material, the coefficients, C and α, density and Poisson’s ratio of which were 0.0102 MPa, 8.04 [49], 0.937 × 10^-3^ g/mm^3^ [39] and 0.475 (nearly incompressible), respectively. The viscous properties of the soft tissue, the relaxation coefficients (g_1_, g_2_) and time constants (τ_1_, τ_2_) were 0.18, 0.12, 0.57 and 6.03, respectively [48]. The outer surface of the foot was modelled as a uniform layer of the skin with a thickness of 1mm [47] and meshed with three-node triangular shell elements. The material properties of the skin were represented as a hyperelastic Ogden material, and coefficients C and α were determined to be 0.122 MPa and 18, respectively [16]. The density and Poisson’s ratio of the skin were assumed to be identical to those of the soft tissue.

**Figure 1.**
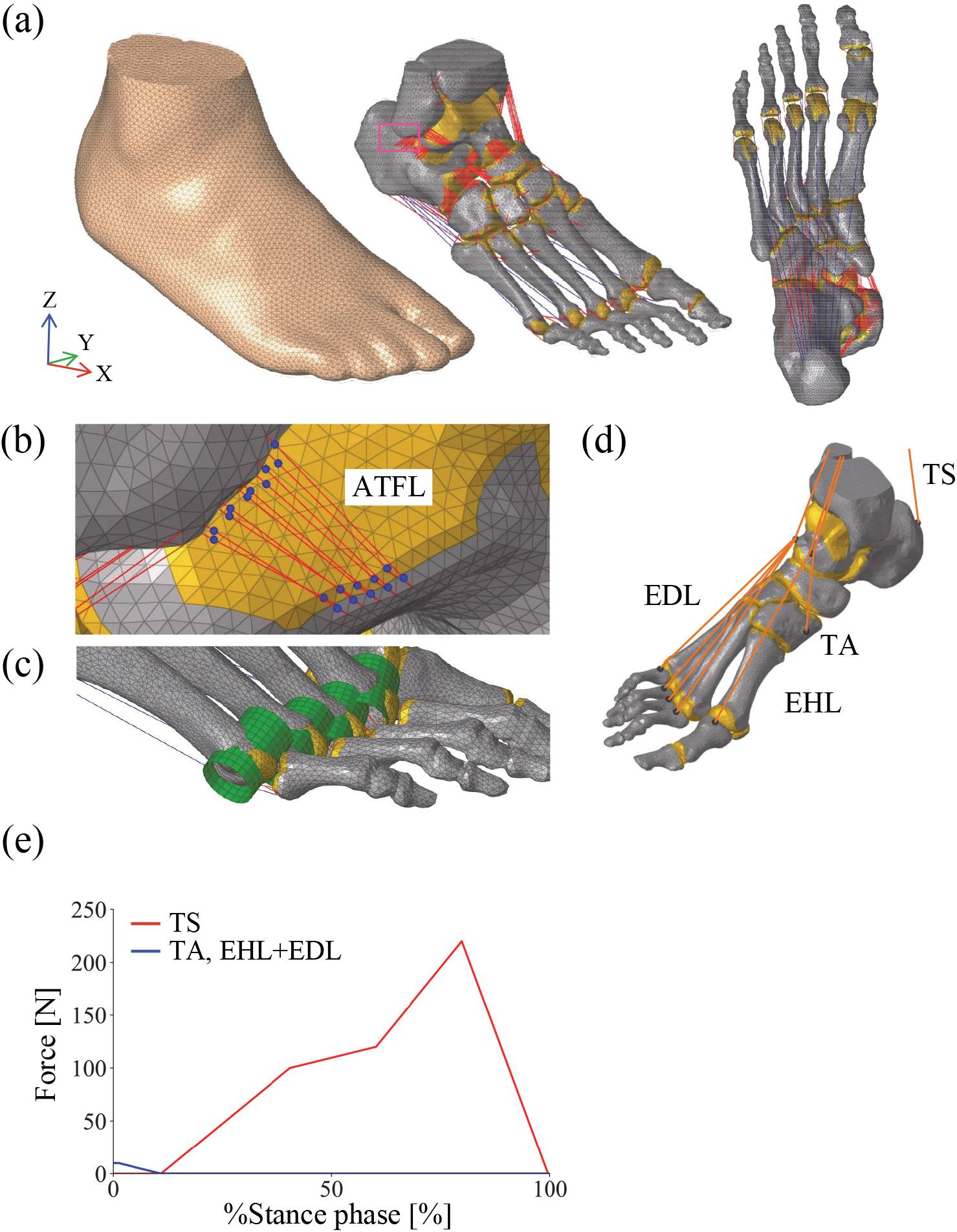
Finite element models of human foot. (a) Skeletal and soft tissue components. Ligaments and the plantar aponeurosis (PA) are shown as red and blue lines, respectively. Cartilages are represented as yellow areas. The dorsal view illustrates the overall configuration of the PA. (b) Anterior talofibular ligament (ATFL). Ligaments are modeled as bundles of an alternating arrangement of tension-only springs and non-penetrating spheres (blue). (c) To prevent penetration of the PA elements during metatarsophalangeal dorsiflexion, the second to fifth metatarsal heads are approximated as cylindrical surfaces (green). (d) Four muscles involved in walking, triceps surae (TS), tibialis anterior (TA), extensor digitorum longus (EDL), and extensor hallucis longus (EHL), are modeled as line segments extending from origins to insertions via intermediate points (orange). (e) Muscle force profiles applied to replicate physiological loading conditions during the stance phase of walking.

Elements corresponding to the articular cavity were manually removed, and thin cartilaginous layers of approximately 0.5 to 1 mm thickness were added on the articular surfaces of the talocrural, talocalcaneal, and calcaneocuboid joints, which were meshed with three-node triangular shell elements. For the other joints, the bone elements corresponding to articular surfaces were selected and assigned as articular cartilage. The Young’s modulus, Poisson’s ratio, and density of the articular cartilage were set to 10 MPa, 0.4, and 0.002 g/mm^3^, respectively [15, 39]. Articular surface-to-surface contact was simulated using a frictionless contact model. The horizontal floor was modeled as a rigid wall, and the contact between the plantar surface of the foot and floor was modeled using a contact model with friction, with the coefficient of static and dynamic friction set to 0.6 [8].

In the foot model outlined in Ito et al. [20], ligaments were depicted as tension-only spring elements linking origin and insertion points according to anatomical atlases [41, 11], resulting in ligament penetration into connecting bones. Moreover, many foot ligaments possess a sheet-like structure with large attachment sites, yet this anatomical feature was not incorporated in the model.

In the present model, each ligament is represented as an alternating arrangement of tension-only springs and spheres, 1.0 mm in diameter, forming a connected series spanning from origin to insertion points (Figure 1b). The spheres do not penetrate the bones, so the ligaments wrapping around bones can be simulated. To accommodate the relatively large attachment sites, ligaments around the talocrural, talocalcaneal, talonavicular, and calcaneocuboid joints are modeled as bundles of 10 such structures each. The Young’s modulus, Poisson’s ratio, and density of the ligament were 260 MPa, 0.4, and 0.001 g/mm^3^ [7]. The cross-sectional areas of the ligaments in the foot were determined as described in Supplementary Information based on Mkandawire et al. [29], Ito et al. [20], Jotoku et al. [22], Siegler et al. [45], Edama et al. [12–14], Taniguchi et al. [51], Won et al. [53], Sarrafian and Kelikian [24], and Maas et al. [40].

Experimentally determining the slack length of ligaments (and other musculoskeletal soft tissue structures) is extremely challenging; therefore, many studies adopt the tissue length in a defined body posture as the slack length [56]. In the present study, the slack length of each ligament was assumed to be 1.1 times its length in the CT-scanned posture. This is because if the slack length is underestimated, the ligaments remain constantly tensioned, resulting in unrealistically high stiffness and contact forces. However, because the ankle and foot were slightly inverted and adducted during scanning, the lateral hindfoot ligaments (e.g., anterior talofibular, calcaneofibular, lateral talocalcaneal, lateral calcaneocuboid, bifurcate, and posterior talofibular ligaments) were likely stretched relative to their slack length. Accordingly, their slack lengths were defined as their lengths in the CT-scanned posture. Similarly, the plantar ligaments (e.g., plantar calcaneocuboid, plantar calcaneonavicular, plantar cubonavicular, plantar cuneonavicular, plantar cuneocuboid, plantar intercuneiform, plantar tarsometatarsal, plantar metatarsal, short plantar, and long plantar ligaments) were not expected to have been shortened during scanning; therefore, their slack lengths were also defined as their lengths in the CT-scanned posture.

The plantar aponeurosis (PA) was modeled using 10 tension-only spring elements that connected the origin and insertion points, with the first PA element passing through the sesamoids beneath the hallux (Figure 1a). To prevent penetration of the second to fifth PA elements during metatarsophalangeal (MP) joint dorsiflexion, the articular surfaces of the metatarsal heads were approximated as cylindrical surfaces, allowing the PA elements to wrap smoothly around them (Figure 1c). The Young’s modulus of the PA was estimated based on Caravaggi et al. [5], who reported that the tensile force of the PA was approximately 1.5 times the body weight when the strain was 0.05. Assuming a body weight of 70 kg and the cross-sectional area of 50 mm^2^ [26], the value was calculated to be 412.02 MPa. The Poisson’s ratio and density of the PAs were 0.4 and 0.001 g/mm^3^, respectively. The slack length of the PA was defined as its length in the CT-scanned posture, as the metatarsophalangeal joints were slightly flexed in that posture.

### Forward dynamic simulation

In the present study, the mechanical interactions of the foot with the ground were simulated via an explicit dynamic analysis. An explicit FE analysis was performed using RADIOSS 2022 (Altair Engineering, Troy, MA, United States). The FUJITSU Supercomputer PRIMEHPC FX1000/Server PRIMERGY GX2570 M6 (Wisteria-O/A) at the Information Technology Center, The University of Tokyo, was used in this study. The time step was set to 1 ms to ensure numerical stability and prevent oscillations.

### Comparisons with cadaveric experiment

The human foot model was validated against a cadaveric study investigating the innate mobility and mechanical properties derived from the anatomy and morphology of the human foot under axial loading [31]. In that experiment, human cadaver feet were subjected to vertical compression without tendon traction, and the resulting three-dimensional movements of the foot bones and tibia were quantified using biplane fluoroscopy [see Figure 1 in Negishi et al. [31]]. The study protocol was approved by the Office for Life Science Research Ethics and Safety at the University of Tokyo (Approval No. 19-295). Clinical trial number: not applicable.

In this experiment, the proximal ends of the tibia and fibula were embedded in a custom-fabricated socket made from a rubber-like polymer and attached to a shaft constrained to translate and rotate only along the vertical axis via a linear guide. We replicated this setup virtually using our developed human foot model. Specifically, the proximal ends of the tibia and fibula were embedded in a rectangular solid made of rubber material (70 mm × 50 mm × 20 mm; Young’s modulus = 4.0 MPa; Poisson’s ratio = 0.475; density = 0.0012 g/mm^3^), which was then fixed to a vertical shaft at the midpoint between the tibia and fibula. The rectangular solid, like the experimental setup, was permitted to move only along and rotate about the vertical axis.

The zero-loading condition was defined as the posture when only the vertical shaft (weighing 3.3 kg) was attached to the model. To reproduce the experimental setup more accurately and avoid introducing unnatural shear forces on the plantar surface, the foot was initially placed heel-first, with the forefoot slightly elevated, before achieving a fully plantigrade posture. This initial positioning was simulated by applying upward forces at the metatarsal heads while axially loading the tibia and fibula. This step was critical to ensure reproducibility, as the initial posture may influence foot mobility under load.

Subsequently, axial loading was applied from the zero-loading condition up to 588 N (equivalent to 60 kg), and the resulting 3D translations and rotations of the foot bones were computed for comparison with the cadaveric results.

To quantify 3D bone kinematics under axial loading, a bone-fixed local coordinate system was established. At the initial position, the three orthogonal axes (x, y, z) of each bone’s local system were aligned with the global axes (X, Y, Z). The X-, Y-, and Z-axes corresponded to anterior, medial and superior directions, respectively. The origin of each bone’s local coordinate system was set at its centroid. Changes in bone position and orientation due to axial loading were described using y–x–z Euler angles, following the method of Ito et al. [18]. These Euler angles captured rotations around the y-, x-, and z-axes, corresponding to plantarflexion–dorsiflexion, inversion–eversion, and internal–external rotation, respectively. Bone-to-bone joint angles for the subtalar, talonavicular, and calcaneocuboid joints were calculated as the motion of the distal bone relative to the proximal bone.

### Static simulation of quiet standing

Physiologically realistic loading condition of the human foot during quiet standing were simulated. Assuming a body mass of 72 kg (the body weight of the CT-scanned subject), a vertical ground reaction force of approximately 353 N was applied to each foot during balanced standing. In addition, based on a structural analysis of the foot during quiet standing [9], the force generated by the triceps surae muscle (via the Achilles tendon) was estimated to be about 50% of the ground reaction force. Therefore, in this study, a downward force of 530 N and an upward force of 177 N were applied to the tibia and the calcaneal tuberosity, respectively. Dynamic calculations were performed as follows: first, an axial force of 353 N was applied to the foot from the initial position without tendon traction, and a converged static solution was obtained. Then, an additional 177 N of axial force was applied to the tibia, along with an upward force at the calcaneal tuberosity, to achieve the final static solution. To validate the human model under quiet standing conditions, we compared the simulated plantar pressure distribution with experimental measurements. The experimental data were obtained from the same subject who underwent CT scanning, using a tactile sensor system (BIG-MAT1300, Nitta, Tokyo, Japan) during balanced standing [20].

### Explicit dynamic simulation of walking

To replicate the dynamic interaction between the foot and the ground during human bipedal walking using the constructed FE model, lower leg kinematics and ground reaction forces (GRFs) were measured from the same subject who underwent CT scanning. The tibia and fibula in the FE foot model were driven by these kinematic data to ensure anatomical consistency in the simulation. Specifically, the motions of markers placed on the medial malleolus, lateral malleolus, and tibial tuberosity were recorded using a motion capture system. After low-pass filtered at a cutoff frequency of 20 Hz, the positions of these three landmarks in the FEM model were matched to the corresponding marker positions measured during walking by minimizing the positional error, and the resulting displacements of the three landmarks were applied to the FEM model as prescribed displacements to drive the explicit dynamic simulation.

However, due to a small discrepancy between the experimental measurements and the model, direct use of the motion capture data did not work. The ground reaction force did not act appropriately when using the height information of the points obtained from motion capture. Therefore, the height data needed to be adjusted so that the vertical component of the ground reaction force would be applied correctly. As an initial step, we reproduced the experimentally observed foot posture and three components of the GRF at 20% of the stance phase by applying external forces to the tibia and fibula. This phase was selected because the tibia is nearly vertical, the plantar surface is fully in contact with the ground, and its angular velocity is relatively low [3, 4, 34]. Additionally, muscle forces from the triceps surae, tibialis anterior, extensor digitorum longus and extensor hallucis longus (Figure 1d) were obtained for the stance phase using inverse dynamics [35] (Figure 1e), and the values at 20% of the stance phase were applied to replicate physiological loading conditions.

From this condition, the experimentally measured displacement of the tibia and the muscle forces were applied in reverse (backward in time) from the 20% stance phase, thereby reconstructing the walking motion up to heel contact. In ideal Newtonian mechanics, the governing equations are time-reversal symmetric, and backward time integration is mathematically valid for purely conservative systems without non-conservative forces (e.g., friction or dissipation). Although our model includes material viscosity and friction, their effects were assumed to be small; thus, backward integration was used to estimate the posture immediately prior to ground contact, which was necessary for calibrating the model’s vertical position. The velocity of the foot model at heel contact was then extracted, reversed, and used as the initial velocity for a forward dynamic simulation. Applying the original tibial displacement data forward in time enabled the reproduction of walking motion in the forward direction.

However, this initial procedure alone did not produce simulated GRFs that matched the experimental data. Therefore, the prescribed tibial displacements were refined. Specifically, the differences between the simulated and experimental GRF components were calculated over time. An eighth-order polynomial function was fitted to each GRF difference waveform using the least squares method, yielding time-dependent correction functions. These functions were scaled by empirically determined coefficients to convert GRF errors into displacement corrections. The resulting adjustments were then added to the original tibial displacement data, and the simulation was rerun. This refined approach led to simulated GRFs that more closely resembled those observed during actual walking.

Three-dimensional bone kinematics during walking were quantified by constructing bone-fixed local coordinate systems, following the approaches of Gutekunst et al. [17] and Negishi et al. [31]. Triaxial joint rotations of the talocrural, talonavicular, and calcaneocuboid joints, as well as the angle between the navicular and first metatarsal, were calculated using y–x–z Euler angles, corresponded to plantarflexion–dorsiflexion, inversion–eversion, and adduction–abduction, respectively. These joint angles were then compared with experimental kinematic data obtained during the stance phase of walking, estimated using a neural network model that tracked foot bone motion based on 41 skin-mounted markers [25]. We also compared the simulated plantar pressure distribution with experimental measurements using the tactile sensor system.

## Results

We compared the translational and rotational responses of foot bones under axial loading in the FE simulation with experimental results from cadaveric feet reported by Negishi et al. [31] (Figure 2). In general, the simulation reproduced the overall trends in bone motion, although some differences in magnitude were observed. Specifically, anterior and inferior translations tended to be slightly greater, while medial displacements were somewhat smaller in the simulation than in the experimental data (Figure 2a). The directions of bone rotations were also consistent between the simulated and measured results (Figure 2b). The mean absolute differences (± standard deviations) in translation between the simulation and experimental data were 2.5 ± 1.4 mm (anteroposterior), 3.1 ± 0.5 mm (mediolateral) and 4.5 ± 1.1 mm (superoinferior). Corresponding angular discrepancies for inversion–eversion, plantarflexion–dorsiflexion, and internal–external rotation were 1.6 ± 0.7°, 2.9 ± 2.3°, and 0.6 ± 0.4°, respectively. Considering that the FE model was not based on the same specimens used in the experimental study, these differences are relatively minor. Furthermore, we compared the simulated plantar pressure distribution during quiet standing with experimental data. The simulated pressure pattern and center of pressure location were largely in agreement with the measured values, while minor discrepancies existed (Figure 3).

**Figure 2.**
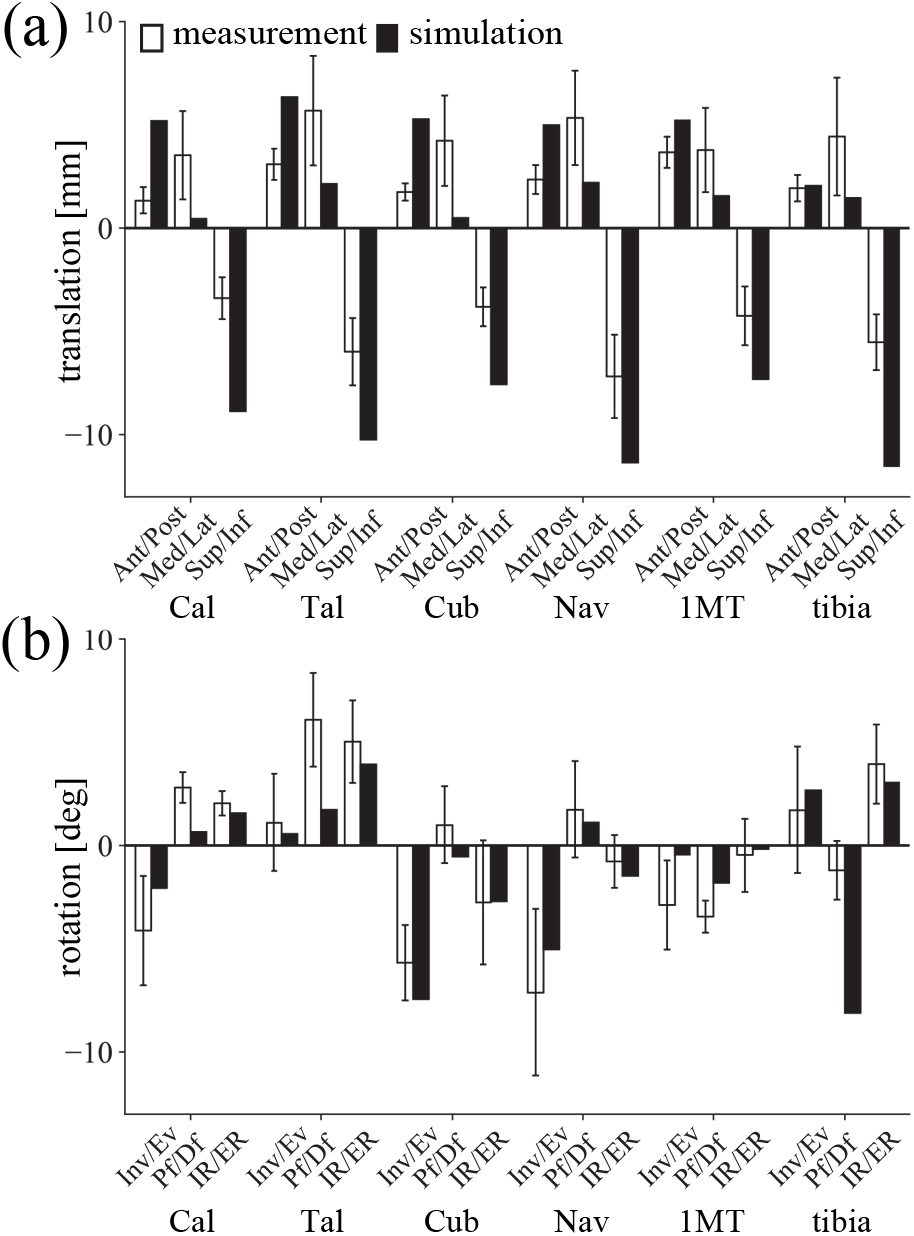
Comparison of simulated foot bone movements under axial loading with experimental data from Negishi et al. [31]. (a) Linear displacements along the anteroposterior, mediolateral, and vertical (superoinferior) axes. Positive values indicate anterior, medial, and superior translation. (b) Rotations within the coronal, sagittal, and transverse planes. Positive values represent inversion, plantarflexion, and internal rotation. Black box: simulation results; White box: experimental data. Abbreviations: Cal = calcaneus; Tal = talus; Cub = cuboid; Nav = navicular; 1MT = first metatarsal.

**Figure 3.**
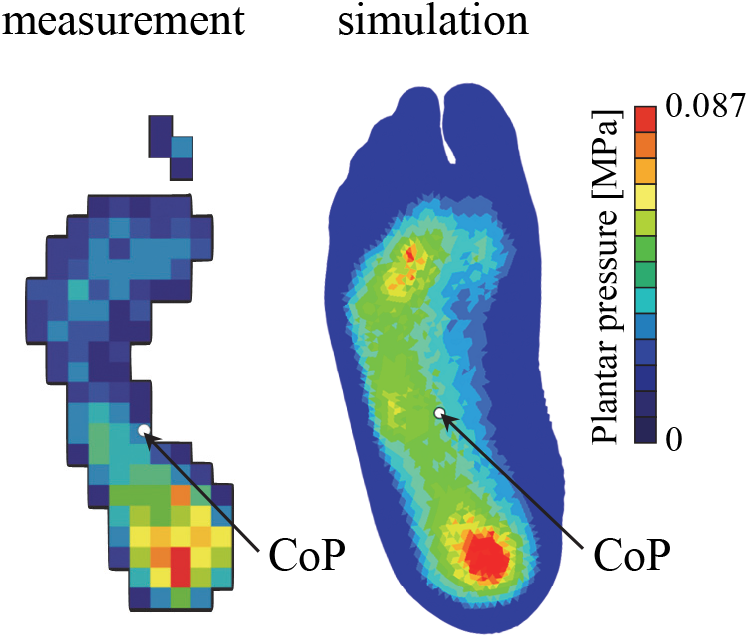
Simulated plantar pressure distribution during quiet standing. The center of pressure (COP) is marked by a white circle. Experimentally measured plantar pressure during quiet standing in a human is shown for comparison.

In the explicit dynamic simulation of walking, the three components of the simulated ground reaction forces during the stance phase were compared with the corresponding experimental measurements (Figure 4a). The simulated ground reaction forces did not perfectly match the measured waveforms; however, key biomechanical characteristics were reasonably reproduced. The anterior-posterior ground reaction force exhibited a braking phase in early stance and a propulsive phase in late stance, with amplitudes closely matching the measured data. The mediolateral component showed a noticeable lateral peak toward the end of stance, which may warrant further investigation, but the overall pattern was largely consistent. The vertical component marginally reproduced the characteristic double-peaked profile observed in human walking. The time-varying plantar pressure distribution obtained from the simulation was compared with the experimentally measured distribution at 10% intervals throughout the stance phase (Figure 4b; however, the results at 20% and 40% were similar to that at 30%, and thus are not shown). Although the patterns were not identical, the center of pressure progressed from the heel to the toes and the overall footprint shape and pressure distribution were generally similar, indicating that the simulated pattern is broadly consistent with the measured data.

**Figure 4.**
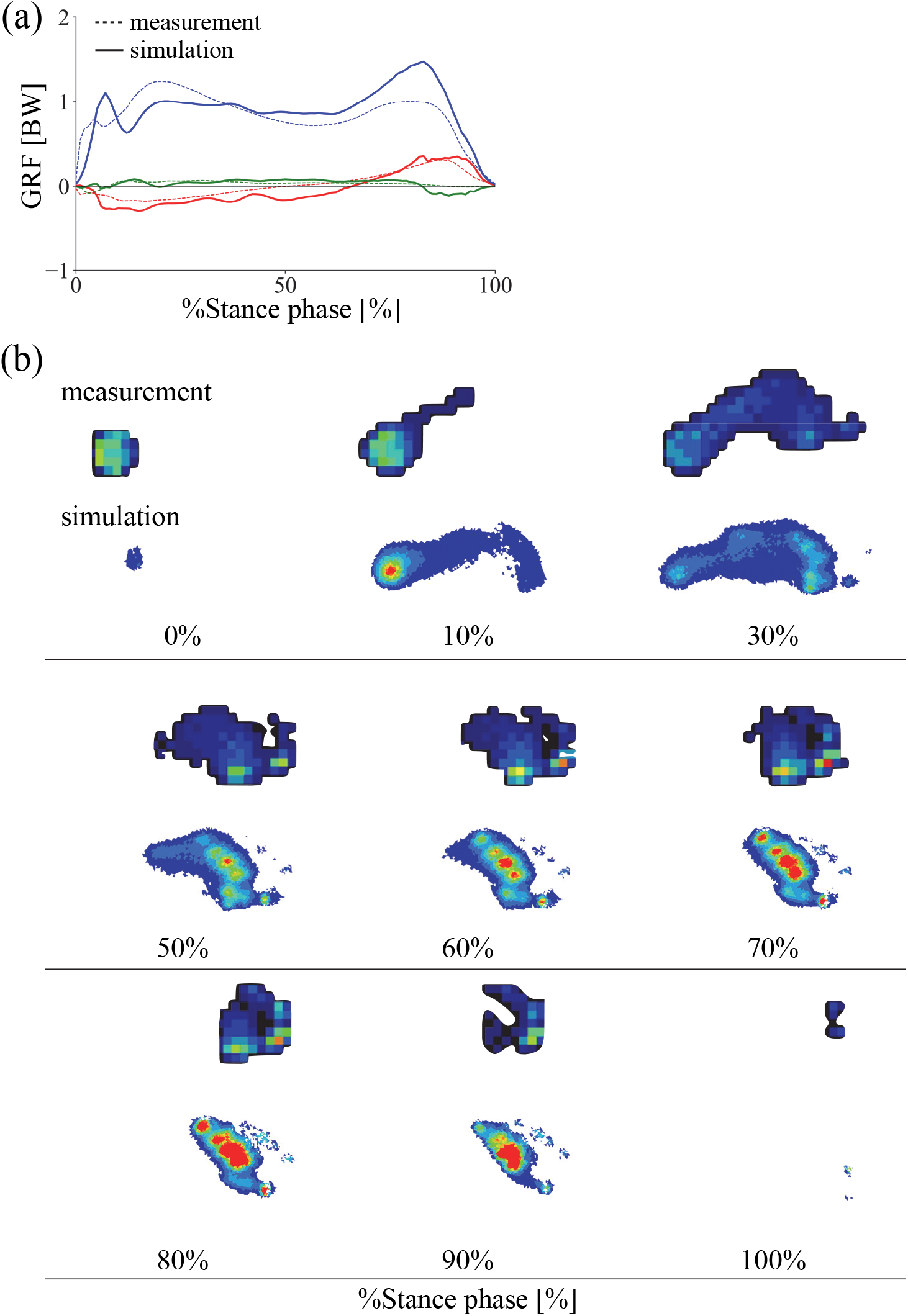
Comparison of simulated and measured ground reaction force (GRF) profiles (a) and plantar pressure distributions (b) during walking. Solid lines represent simulation results; dashed lines represent experimental data. Red = anteroposterior GRF; Green = mediolateral GRF; Blue = vertical GRF. Positive values indicate posterior, medial, and superior directions.

The simulated foot movements during the stance phase were presented in Figure 5 (See Supplementary Information for a video), and the changes in joint angles and their comparisons with the experimentally obtained corresponding joint angles estimated using a neural network model that tracked foot bone motion based on 41 skin-mounted markers [25] were presented in Figure 6a. Although there are some discrepancies, the changes in foot joint angles in the simulation are generally consistent with the experimental kinematic data. In addition, the maximum and minimum values of the estimated joint angle profiles and the timing of these peaks generally aligned with those reported in previous studies using bi-planar fluoroscopy [23, 38], indicating that our simulation is also largely consistent with those measured data.

**Figure 5.**
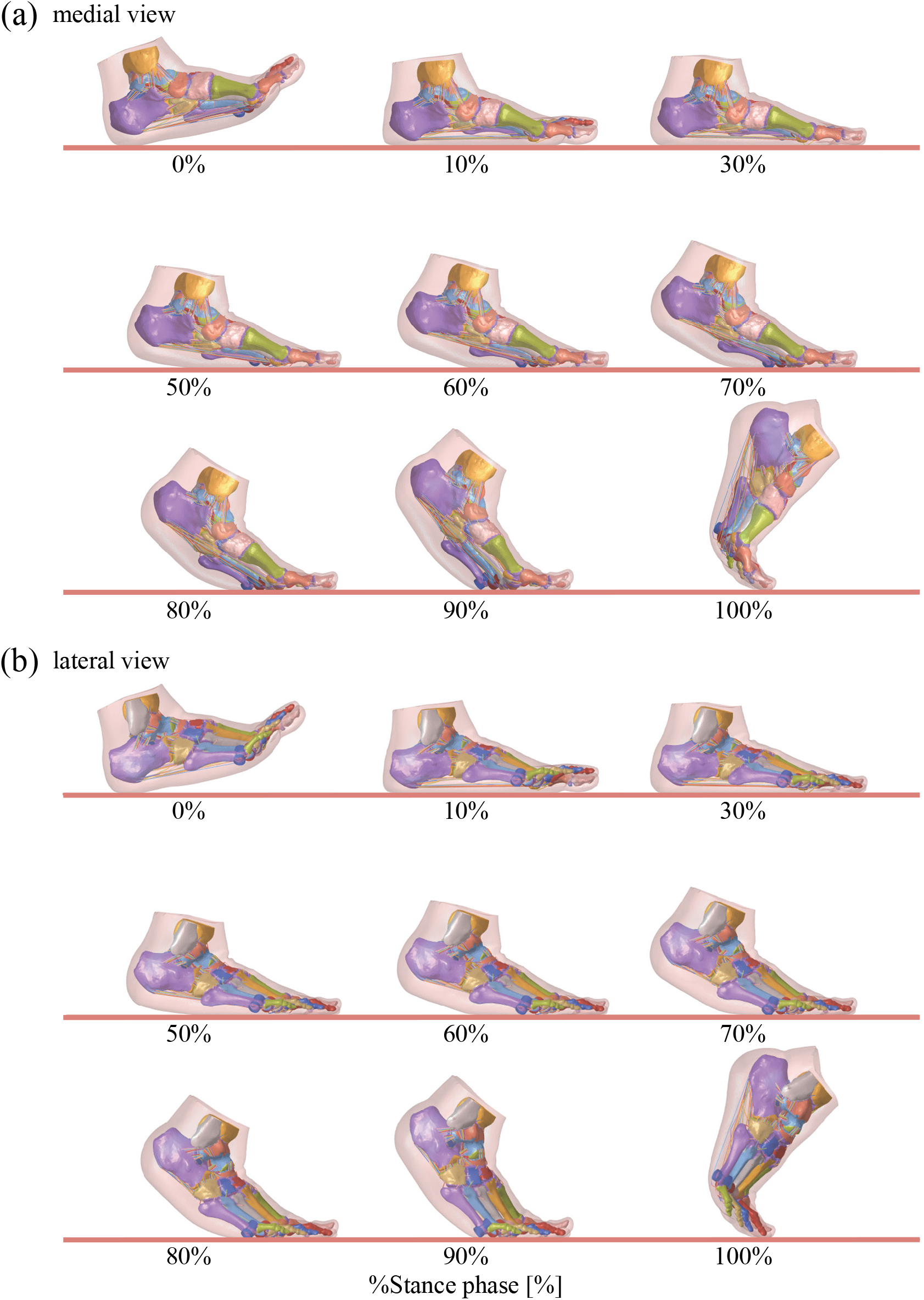
Simulated foot movements during walking. (a) Medial and (b) lateral views. The medial view is mirrored to match the left-to-right progression of time.

**Figure 6.**
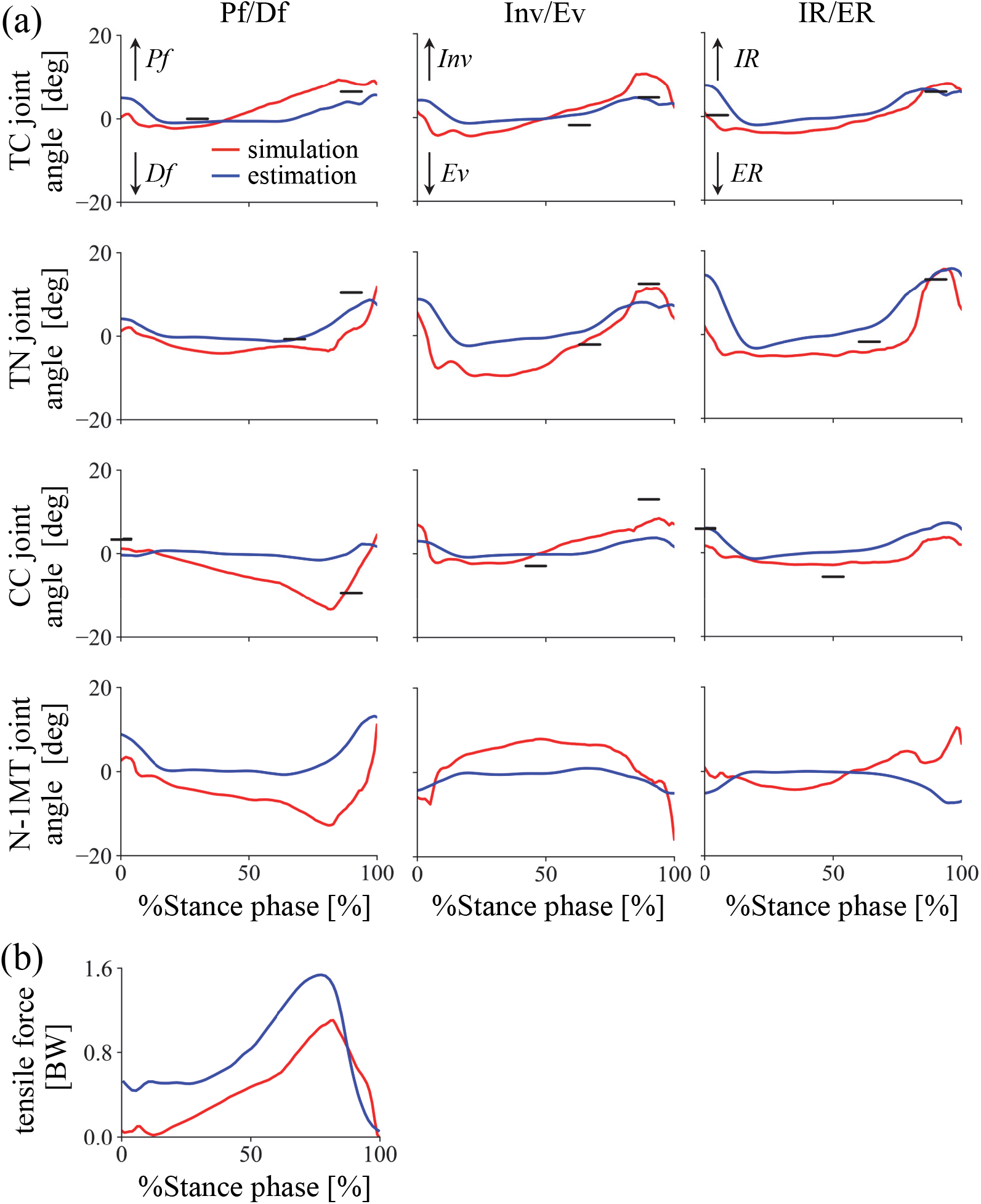
Comparison of simulated joint angles (a) and plantar aponeurosis force profiles (b) during walking. Simulated joint angles include the talocalcaneal (TC), talonavicular (TN), and calcaneocuboid (CC) joints, as well as the rotational angle of the first metatarsal relative to the navicular (N–1MT). Red lines: simulation; Blue lines: estimates based on experimental data from Matsumoto et al. [25] and Caravaggi et al. [5]. Horizontal bars indicate the mean peak angles and their timing as measured using biplanar fluoroscopy based on published data (Koo et al., [23]; Phan et al., [38]).

The simulated tension profile of the PA during walking (Figure 6b) showed a pattern of increasing force throughout most of the stance phase, consistent with the findings of Caravaggi et al. [5, 6], who reported that the tensile force of the PA reached approximately 1.5 times body weight. In our simulation, the peak tensile force reached approximately 1.1 times body weight, showing a similar waveform and generally good agreement with their results.

Figures 7 and 8 illustrate the time-varying von Mises stress distribution in the bones and soft tissues, respectively, during walking (See Supplementary Information for a video). Although such stress changes cannot be measured directly in vivo, the simulation enables their visualization. The results suggest that bone stress is concentrated in the metatarsals, while soft tissue stress is primarily distributed under the plantar regions of the calcaneus and metatarsophalangeal joints.

**Figure 7.**
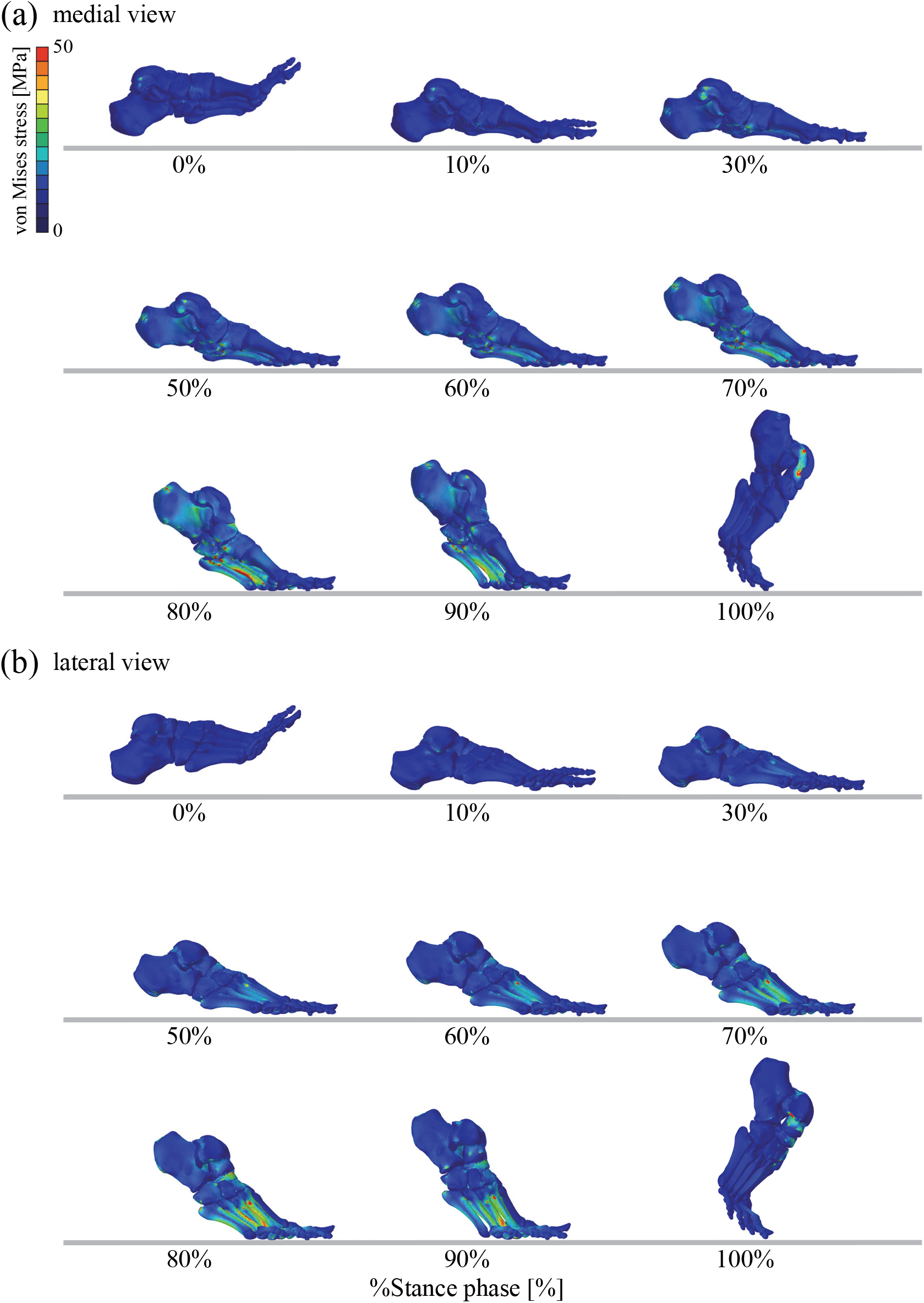
Simulated von Mises stress distributions of the foot bones during walking. (a) Medial and (b) lateral views. The medial view is mirrored to match the left-to-right progression of time.

**Figure 8.**
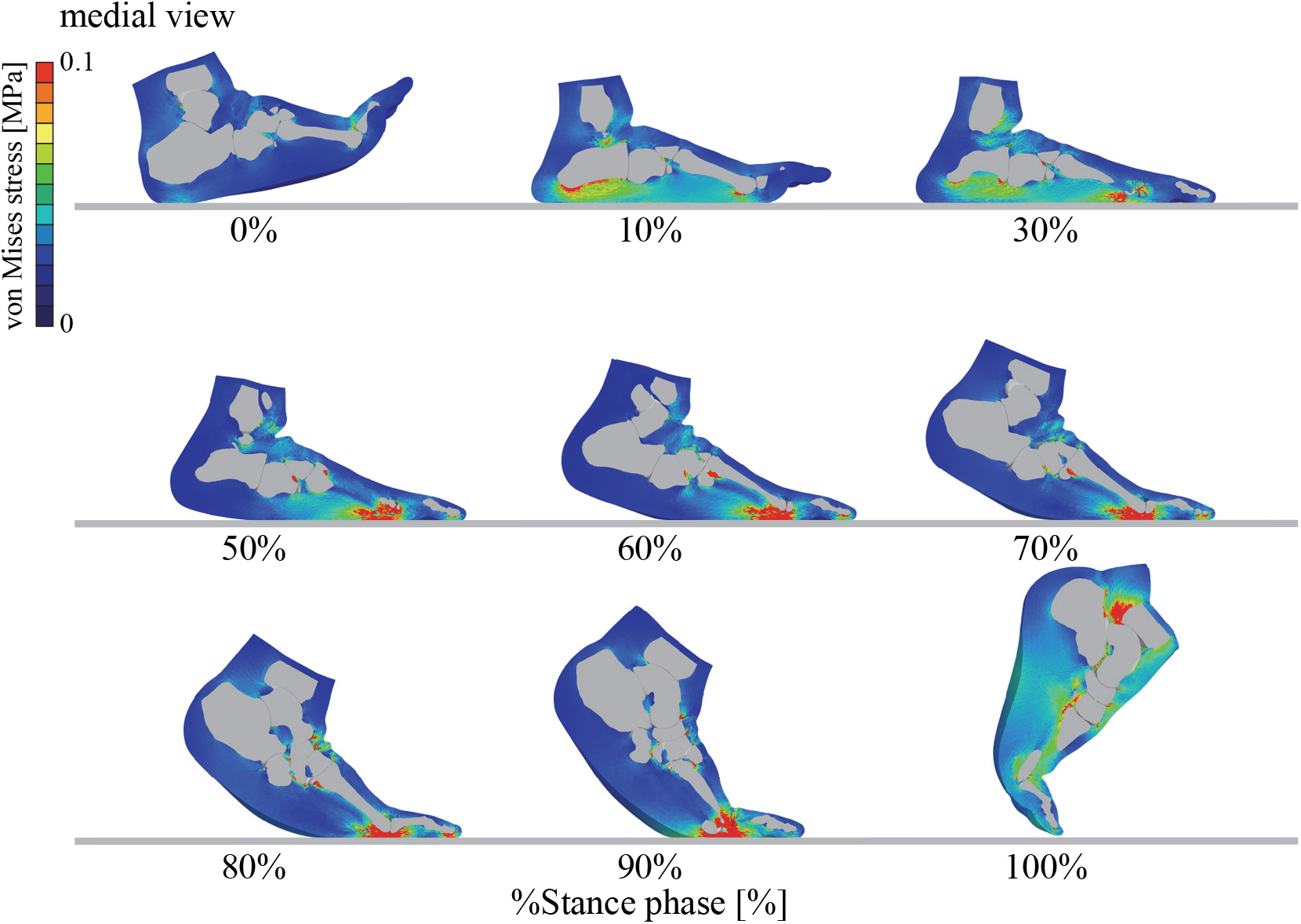
Simulated von Mises stress distributions of the soft tissue during walking.

## Discussion

This study demonstrated that the explicit finite element simulation of the foot, driven by experimentally measured tibial kinematics and estimated muscle forces, successfully replicated key mechanical behaviors of the foot as it interacts with the ground during walking. The simulation results showed reasonable agreement with experimental data in terms of kinematics, ground reaction forces, and plantar pressure distribution throughout the stance phase. Considering that the model foot and the foot used for the kinematic measurements (Figures 2 and 6) were not identical, perfect agreement could not be expected; nevertheless, the correspondence observed was reasonably consistent. This level of agreement indicates that the model is sufficiently accurate for qualitative analysis and hypothesis testing, although refinements in geometry and subject-specific material properties would be necessary for precise quantitative predictions. These findings suggest that the model captures essential aspects of foot mechanics during walking. Although some discrepancies were observed likely due to simplifications in the modeling of soft tissues, ligaments, and contact properties, the model provides a valuable framework that can be applied to investigate the relationship between form and function in the musculoskeletal system, as well as to advance our understanding of pathogenesis of foot and ankle disorders.

By enabling the estimation of internal forces, stresses, and strains within foot structures during gait, parameters that are not directly measurable in experimental settings, this model offers valuable insights into the biomechanics of bones, soft tissues, ligaments, and joints during human walking. Although the present results require further validation for quantitative accuracy, they are qualitatively reasonable and align with known biomechanical patterns. Such information is essential for understanding the development of foot pathologies, including plantar fasciitis [27, 28], hallux valgus [33], ankle osteoarthritis [42, 43], and diabetic foot ulcers [19]. Moreover, this modeling framework can be extended to assess the mechanical effects of footwear design, orthotic interventions, and surgical procedures.

The adoption of an explicit dynamic simulation approach proved advantageous for handling complex contact interactions and large deformations within the foot during walking. It must be noted, however, that the method required substantial computational time. In the present study, the calculation of foot dynamics from foot contact to toe-off (Figure 5) required approximately 14 hours using the supercomputer. While such computational times are acceptable for research purposes, clinical or design applications would require significant reduction in runtime through model optimization, parallel computing, or reduced-order modeling techniques. Balancing computational efficiency with anatomical and biomechanical realism remains a challenge for applying such simulations to the detailed understanding of musculoskeletal dynamics and their medical applications.

This study has several limitations. First, the foot model for each species was based on CT data from a single individual and therefore does not capture the natural variation in foot morphology. Second, the material properties used in the finite element analysis were compiled from various literature sources. As a result, parameters such as ligament cross-sectional areas were not individually scaled due to insufficient data for accurate adjustment. Additionally, the foot’s soft tissues were simplified as a single homogeneous material, whereas in reality they consist of structurally and mechanically distinct components, including fat, muscle, and tendons. Although the soft tissue material parameter was obtained from experimental measurements of the heel pad as a whole, rather than separately for individual tissue components, making this simplification reasonable, the anatomical differences between the modeled and actual foot likely contributed to discrepancies in the simulation results. Despite these limitations, however, we believe that the present simulation successfully captured key aspects of foot mechanics and offers meaningful insights into foot function, highlighting the potential of this modeling approach for future research in comparative morphology, biomechanics, and clinical applications. Therefore, the present study contributes to a deeper understanding of the mechanical principles underlying foot function in both evolutionary and clinical contexts.

## Author contributions

NO conceived and designed the study and supervised the project; KI and YM performed the simulation and data analysis; NO, HS and TN evaluated the simulation; NO drafted the manuscript, and all authors edited and approved the manuscript prior to submission.

## Data availability

The datasets generated and/or analyzed during the current study are available from the corresponding author upon reasonable request.

## References

1. Akrami M, Qian Z, Zou Z, Howard D, Nester CJ, Ren L. Subject-specific finite element modelling of the human foot complex during walking: sensitivity analysis of material properties, boundary and loading conditions. Biomech Model Mechanobiol. 2018; 10.1007/s10237-017-0978-3

2. Aoi S, Ogihara N, Funato T, Tsuchiya K. Sensory regulation of stance-to-swing transition in generation of adaptive human walking: A simulation study. Robotics Auton Syst. 2012; 10.1016/j.robot.2011.12.005

3. Bianchi L, Angelini D, Orani GP, Lacquaniti F. Kinematic coordination in human gait: relation to mechanical energy cost. J Neurophysiol. 1998; 10.1152/jn.1998.79.4.2155

4. Borghese NA, Bianchi L, Lacquaniti F. Kinematic determinants of human locomotion. J Physiol Lond. 1996; 10.1113/jphysiol.1996.sp021539

5. Caravaggi P, Pataky T, Goulermas JY, Savage R, Crompton R. A dynamic model of the windlass mechanism of the foot: evidence for early stance phase preloading of the plantar aponeurosis. J Exp Biol. 2009; 10.1242/jeb.025767

6. Caravaggi P, Pataky T, Gunther M, Savage R, Crompton R. Dynamics of longitudinal arch support in relation to walking speed: contribution of the plantar aponeurosis. J Anat. 2010; 10.1111/j.1469-7580.2010.01261.x

7. Chen TLW, Wong DWC, Wang Y, Lin J, Zhang M. Foot arch deformation and plantar fascia loading during running with rearfoot strike and forefoot strike: A dynamic finite element analysis. J Biomech. 2019; 10.1016/j.jbiomech.2018.12.007

8. Chen WM, Lee T, Lee PV, Lee JW, Lee SJ. Effects of internal stress concentrations in plantar soft-tissue--A preliminary three-dimensional finite element analysis. Med Eng Phys. 2010; 10.1016/j.medengphy.2010.01.001

9. Cheung JTM, Zhang M, Leung AKL, Fan YB. Three-dimensional finite element analysis of the foot during standing--a material sensitivity study. J Biomech. 2005; 10.1016/j.jbiomech.2004.05.035

10. Di Russo A, Stanev D, Armand S, Ijspeert A. Sensory modulation of gait characteristics in human locomotion: A neuromusculoskeletal modeling study. PLoS Comput Biol. 2021; 10.1371/journal.pcbi.1008594

11. Drake RL, Vogl AW, Mitchell AWM, Tibbitts RM, Richardson PE. Gray’s Atlas of Anatomy. London: Churchill Livingstone; 2008.

12. Edama M, Ikezu M, Kaneko F, Kikumoto T, Takabayashi T, Hirabayashi R, Inai T, Kageyama I. Morphological features of the bifurcated ligament. Surg Radiol Anat. 2019; 10.1007/s00276-018-2089-y

13. Edama M, Takabayashi T, Hirabayashi R, Yokota H, Sekine C, Inai T, Matsuzawa K, Otsuki T, Maruyama S, Kageyama I. Morphological features of the lateral plantar ligament of the transverse metatarsal arch. Clin Anat. 2021; 10.1002/ca.23687

14. Edama M, Takabayashi T, Inai T, Hirabayashi R, Ikezu M, Kaneko F, Matsuzawa K, Kageyama I. Morphological features of the cervical ligament. Surg Radiol Anat. 2020; 10.1007/s00276-019-02364-y

15. Gefen A. Plantar soft tissue loading under the medial metatarsals in the standing diabetic foot. Med Eng Phys. 2003; 10.1016/s1350-4533(03)00029-8

16. Gu Y, Li J, Ren X, Lake MJ, Zeng Y. Heel skin stiffness effect on the hind foot biomechanics during heel strike. Skin Res Technol. 2010; 10.1111/j.1600-0846.2010.00425.x

17. Gutekunst DJ, Liu L, Ju T, Prior FW, Sinacore DR. Reliability of clinically relevant 3D foot bone angles from quantitative computed tomography. J Foot Ankle Res. 2013; 10.1186/1757-1146-6-38

18. Ito K, Hosoda K, Shimizu M, Ikemoto S, Nagura T, Seki H, Kitashiro M, Imanishi N, Aiso S, Jinzaki M, Ogihara N. Three-dimensional innate mobility of the human foot bones under axial loading using biplane X-ray fluoroscopy. R Soc Open Sci. 2017; 10.1098/rsos.171086

19. Ito K, Maeda K, Fujiwara I, Hosoda K, Nagura T, Lee T, Ogihara N. Dynamic measurement of surface strain distribution on the foot during walking. J Mech Behav Biomed Mater. 2017; 10.1016/j.jmbbm.2016.12.009

20. Ito K, Nakamura T, Suzuki R, Negishi T, Oishi M, Nagura T, Jinzaki M, Ogihara N. Comparative functional morphology of human and chimpanzee feet based on three-dimensional finite element analysis. Front Bioeng Biotechnol. 2021; 10.3389/fbioe.2021.760486

21. Jo S, Massaquoi SG. A model of cerebrocerebello-spinomuscular interaction in the sagittal control of human walking. Biol Cybern. 2007; 10.1007/s00422-006-0126-0

22. Jotoku T, Kinoshita M, Okuda R, Abe M. Anatomy of ligamentous structures in the tarsal sinus and canal. Foot Ankle Int. 2006; 10.1177/107110070602700709

23. Koo S, Lee KM, Cha YJ. Plantar-flexion of the ankle joint complex in terminal stance is initiated by subtalar plantar-flexion: A bi-planar fluoroscopy study. Gait Posture 2015; 10.1016/j.gaitpost.2015.07.009

24. Maas NM, van der Grinten M, Bramer WM, Kleinrensink GJ. Metatarsophalangeal joint stability: a systematic review on the plantar plate of the lesser toes. J Foot Ankle Res. 2016; 10.1186/s13047-016-0165-2

25. Matsumoto Y, Hakukawa S, Seki H, Nagura T, Imanishi N, Jinzaki M, Kanemura N, Ogihara N. Estimating three-dimensional foot bone kinematics from skin markers using a deep learning neural network model. J Biomech. 2024; 10.1016/j.jbiomech.2024.112252

26. Matsumoto Y, Ogihara N. Direct visualization and measurement of the plantar aponeurosis behavior in foot arch deformation via the windlass mechanism. Clin Anat. 2025; 10.1002/ca.24171

27. Matsumoto Y, Ogihara N, Hanawa H, Kokubun T, Kanemura N. Novel multi-segment foot model incorporating plantar aponeurosis for detailed kinematic and kinetic analyses of the foot with application to gait studies. Front Bioeng Biotechnol. 2022; 10.3389/fbioe.2022.894731

28. Matsumoto Y, Ogihara N, Kosuge S, Hanawa H, Kokubun T, Kanemura N. Sex differences in the kinematics and kinetics of the foot and plantar aponeurosis during drop-jump. Sci Rep. 2023; 10.1038/s41598-023-39682-6

29. Mkandawire C, Ledoux WR, Sangeorzan BJ, Ching RP. Foot and ankle ligament morphometry. J Rehabil Res Dev. 2005; 10.1682/jrrd.2004.08.0094

30. Mo FH, Li YD, Li JJ, Zhou SY, Yang ZR. A three-dimensional finite element foot-ankle model and its personalisation methods analysis. Int J Mech Sci. 2022; 10.1016/j.ijmecsci.2022.107108

31. Negishi T, Ito K, Hosoda K, Nagura T, Ota T, Imanishi N, Jinzaki M, Oishi M, Ogihara N. Comparative radiographic analysis of three-dimensional innate mobility of the foot bones under axial loading of humans and African great apes. R Soc Open Sci. 2021; 10.1098/rsos.211344

32. Neptune RR, Clark DJ, Kautz SA. Modular control of human walking: a simulation study. J Biomech. 2009; 10.1016/j.jbiomech.2009.03.009

33. Nozaki S, Watanabe K, Katayose M, Yamatsu K, Teramoto A, Ogihara N. Three-dimensional morphological variations in the calcaneus and talus in relation to the hallux valgus angle. Ann Anat, 2023; 10.1016/j.aanat.2023.152053

34. Ogihara N, Kikuchi T, Ishiguro Y, Makishima H, Nakatsukasa M. Planar covariation of limb elevation angles during bipedal walking in the Japanese macaque. J R Soc Interface 2012; 10.1098/rsif.2012.0026

35. Ogihara N, Yamazaki N. Generation of human bipedal locomotion by a bio-mimetic neuro-musculo-skeletal model. Biol Cybern. 2001; 10.1007/PL00007977

36. Oku H, Ide N, Ogihara N. Forward dynamic simulation of Japanese macaque bipedal locomotion demonstrates better energetic economy in a virtualised plantigrade posture. Commun Biol. 2021; 10.1038/s42003-021-01831-w

37. Pandy MG, Andriacchi TP. Muscle and joint function in human locomotion. Annu Rev Biomed Eng. 2010; 10.1146/annurev-bioeng-070909-105259

38. Phan CB, Shin G, Lee KM, Koo S. Skeletal kinematics of the midtarsal joint during walking: Midtarsal joint locking revisited. J Biomech. 2019; 10.1016/j.jbiomech.2019.07.031

39. Qian Z, Ren L, Ding Y, Hutchinson JR, Ren L. A dynamic finite element analysis of human foot complex in the sagittal plane during level walking. PLoS One 2013; 10.1371/journal.pone.007942

40. Sarrafian SK, Kelikian AS. Sarrafian’s anatomy of the foot and ankle: descriptive, topographic, functional. Philadelphia: Wolters Kluwer Health/Lippincott Williams & Wilkins; 2011.

41. Schuenke M, Schulte E, Schumacher U. Atlas of Anatomy: General Anatomy and Musculoskeletal System. New York: Thieme; 2006.

42. Seki H, Nozaki S, Ogihara N, Kokubo T, Nagura T. Morphological features of the non-affected side of the hindfoot in patients with unilateral varus ankle osteoarthritis. Ann Anat. 2024; 10.1016/j.aanat.2023.152198

43. Seki H, Ogihara N, Kokubo T, Suda Y, Ishii K, Nagura T. Visualization and quantification of the degenerative pattern of the talus in unilateral varus ankle osteoarthritis. Sci Rep. 2019; 10.1038/s41598-019-53746-6

44. Sellers WI, Dennis LA, Crompton RH. Predicting the metabolic energy costs of bipedalism using evolutionary robotics. J Exp Biol. 2003; 10.1242/jeb.00205

45. Siegler S, Block J, Schneck CD. The mechanical characteristics of the collateral ligaments of the human ankle joint. Foot Ankle 1988; 10.1177/107110078800800502

46. Song S, Geyer H. A neural circuitry that emphasizes spinal feedback generates diverse behaviours of human locomotion. J Physiol. 2015; 10.1113/JP270228

47. Strzalkowski ND, Triano JJ, Lam CK, Templeton CA, Bent LR. Thresholds of skin sensitivity are partially influenced by mechanical properties of the skin on the foot sole. Physiol Rep. 2015; 10.14814/phy2.12425

48. Suzuki R, Ito K, Lee T, Ogihara N. In-vivo viscous properties of the heel pad by stress-relaxation experiment based on a spherical indentation. Med Eng Phys. 2017; 10.1016/j.medengphy.2017.10.010

49. Suzuki R, Ito K, Lee T, Ogihara N. Parameter identification of hyperelastic material properties of the heel pad based on an analytical contact mechanics model of a spherical indentation. J Mech Behav Biomed Mater. 2017; 10.1016/j.jmbbm.2016.09.027

50. Taga G. A model of the neuro-musculo-skeletal system for human locomotion. I. Emergence of basic gait. Biol Cybern. 1995; 10.1007/BF00204048

51. Taniguchi A, Tanaka Y, Takakura Y, Kadono K, Maeda M, Yamamoto H. Anatomy of the spring ligament. J Bone Joint Surg Am. 2003; 10.2106/00004623-200311000-00018

52. Wang Y, Wong DW, Tan Q, Li Z, Zhang M. Total ankle arthroplasty and ankle arthrodesis affect the biomechanics of the inner foot differently. Sci Rep. 2019; 10.1038/s41598-019-50091-6

53. Won HJ, Won HS, Oh CS, Han SH, Chung IH, Suh JS, Lee WC. Posterior tibiotalar ligament: an anatomic study correlated with MRI. Clin Anat. 2014; 10.1002/ca.22302

54. Yamazaki N, Hase K, Ogihara N, Hayamizu N. Biomechanical analysis of the development of human bipedal walking by a neuro-musculo-skeletal model. Folia Primatol (Basel). 1996; 10.1159/000157199

55. Zajac FE, Neptune RR, Kautz SA. Biomechanics and muscle coordination of human walking: part II: lessons from dynamical simulations and clinical implications. Gait Posture 2003; 10.1016/s0966-6362(02)00069-3

56. Zhang Q, Adam NC, Hosseini Nasab SH, Taylor WR, Smith CR. Techniques for in vivo measurement of ligament and tendon strain: A review. Ann Biomed Eng. 2021; 10.1007/s10439-020-02635-5

