## supplementary information for "Simulating human foot mechanics during walking based on an anatomically detailed forward dynamic finite element model"

Corresponding Author

Naomichi Ogihara

Department of Biological Science, Graduate School of Science,

The University of Tokyo,

7-3-1 Hongo, Bukyo-ku, Tokyo 113-0033, Japan

**Supplementary material**

**Supplementary table**

Cross-sectional areas of the ligaments and corresponding references (cross-sectional area.xlsx).

**Supplementary video**

Simulated temporal changes in foot movements and von Mises stress distributions of the foot bones and soft tissue during walking (FEM_walking.mp4). The video sequentially presents the simulated foot motion, the von Mises stress distributions of the foot bones (shown from medial and lateral views, respectively), and the von Mises stress distribution within the soft tissue. The medial view is mirrored to match the left-to-right temporal progression of the walking cycle.
